# Automatic Identification and Avoidance of Axon Bundle Activation for Epiretinal Prosthesis

**DOI:** 10.1101/2021.02.03.429317

**Authors:** Pulkit Tandon, Nandita Bhaskhar, Nishal Shah, Sasi Madugula, Lauren Grosberg, Victoria H. Fan, Pawel Hottowy, Alexander Sher, Alan M. Litke, E.J. Chichilnisky, Subhasish Mitra

## Abstract

Retinal prostheses must be able to activate cells in a selective way in order to restore high-fidelity vision. However, inadvertent activation of far-away retinal ganglion cells (RGCs) through electrical stimulation of axon bundles can produce irregular and poorly controlled percepts, limiting artificial vision. Therefore, the problem of *axon bundle activation* can be defined as the axonal stimulation of RGCs with unknown soma and receptive field locations, typically outside the electrode array. Here, a new algorithm is presented that utilizes electrical recordings to determine the stimulation current amplitudes above which bundle activation occurs. The method exploits several spatiotemporal characteristics of electrically-evoked spikes to overcome the challenge of detecting small axonal spikes in extracellular recordings. The algorithm was validated using large-scale *ex vivo* stimulation and recording experiments in macaque retina, by comparing algorithmically and manually identified bundle activation thresholds. The algorithm could be used in a closed-loop manner by a future epiretinal prosthesis to reduce poorly controlled visual percepts associated with bundle activation. The method may also be applicable to other types of retinal implants and to cortical implants.

**Contributions:** PT developed the algorithm and analyzed the data, with input from SMi and EJC. NB and NS helped with the analysis. SMa and LG performed dissections and collected the data. PT and VFH performed manual identification. PH, AS and AML developed and supported recording hardware and software. PT, EJC and SMi wrote the manuscript. NS and SMa edited it. EJC and SMi supervised the project.

## Introduction

Retinal prostheses are designed to restore partial vision in patients with photoreceptor degenerative diseases such as age-related macular degeneration and retinitis pigmentosa. These devices aim to overcome the loss of photoreceptors by electrically stimulating the downstream retinal circuitry through current injection via multi-electrode arrays (MEAs) [1,2]. In an *epiretinal prosthesis*, the MEA is placed on the anterior surface of the retina in order to precisely stimulate retinal ganglion cells (RGCs), ideally with single-cell resolution, to emulate naturally-evoked visual perception [2–7]. However, a major challenge in achieving this goal is inadvertent electrical activation of the numerous RGC axons in the nerve fiber layer between the electrodes and RGCs. Activation of axons has been shown to produce irregular arc-shaped phosphenes in patients with epiretinal implants, distorting their artificial visual perception [8–10]. Hence, avoiding indiscriminate axon bundle stimulation [4] could drastically improve artificial vision.

A bidirectional retinal prosthesis (i.e. one with both read and write capability) could substantially enhance the ability to detect and avoid undesired visual percepts. At present, retinal implants rely on patient feedback to determine the visual percepts elicited by electrical stimulation [8,9], but this approach would be prohibitively time-consuming for a prosthesis based on large-scale MEAs [9–11]. To determine the artificial visual signal produced by electrical stimulation, an ideal epiretinal prosthesis would not only stimulate but also record the electrically-evoked and spontaneous spiking activity of each RGC over the MEA, identify its location and cell type, and thus estimate its expected contribution to visual perception [5,12–15]. However, if the soma of a cell lies off the array, its location cannot be identified, and hence the spatial contribution to visual perception introduced by stimulating the cell is uncertain. Thus, the problem of *axon bundle activation* is defined as the activation of off-array cells, and the *axon bundle threshold* for each stimulating electrode is defined as the lowest current amplitude at which the activity in any off-array cell is observed [4] (Figure 1). Although it is possible that unrecorded on-array cells also contribute to unaccounted visual percepts, this work assumes that the high-density MEA allows detection and identification of the majority of the on-array cells [13,16–19].

**Figure 1.**
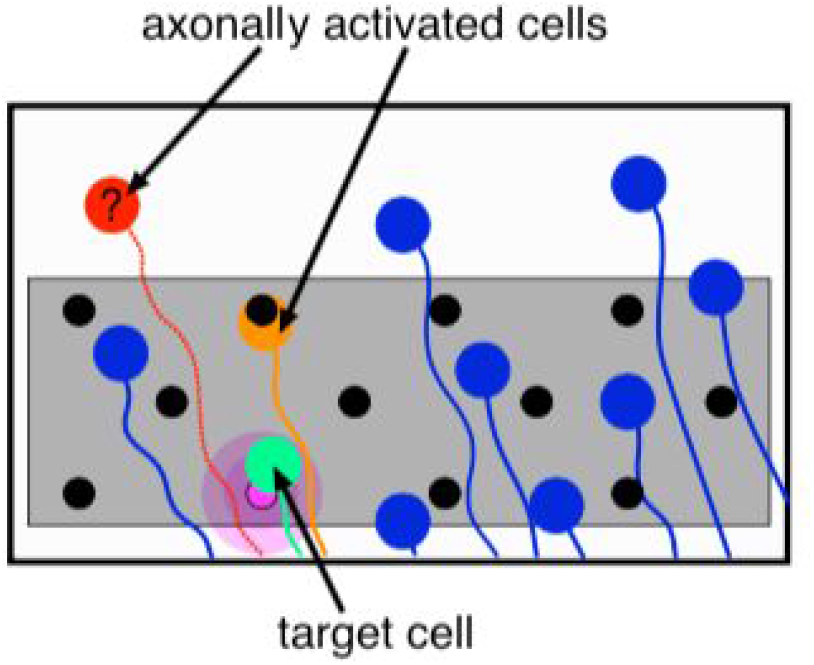
Axon bundle activation. The MEA (gray shaded region) and electrodes (black dots) partially cover the area of the retina containing retinal ganglion cells (RGCs) (large colored dots) with axons (curves) that course to the optic nerve. To activate a target RGC (green), current is typically passed through an electrode near it (pink dot). But this can lead to activating bypassing axons near the stimulating electrode (pink patch). If an axonal spike is evoked in a RGC with its soma on the array (orange), then its location and contribution to artificial vision can be determined. But if the soma of the activated RGC lies off the array (red), then the cell cannot be located and its contribution to vision is unknown.

The current state-of-the-art method for detecting axon bundle activation involves manually analyzing post-stimulation MEA recordings (see Results), which can take days for a single recording with hundreds of electrodes. This is not a feasible approach in a prosthetic device that could require frequent recalibration to ensure reliable performance. Other methods proposed in the literature to detect axonal activation utilize optical recording or examine isolated neurons *in vitro* [20–28]. However, these methods cannot be used in an *in vivo* implant. One closely related study used heuristics based on hand-crafted features for which computation grows superlinearly in the number of recorded electrodes [4]. Moreover, it does not necessarily discriminate on-array axonal activation from off-array activation, leading to certain stimulation levels being classified as above axon bundle threshold even though the elicited visual percepts can be determined (e.g. Figure 1, stimulation of orange but not red RGC). This may indicate fewer allowable stimulation levels without axon bundle activation, and thus may limit the utility of the implant. Alternative techniques to avoid axonal activation such as varying the pulse duration or frequency have also been proposed [29]. Even so, use of these techniques does not eliminate all off-array activation, and an algorithm to detect axon bundle threshold remains important.

Here, a simple and principled algorithm is presented to determine axon bundle thresholds. The algorithm was applied to detect bundle activation in *ex vivo* electrical stimulation and recording data from peripheral macaque retina, and the results suggest that the algorithm is accurate and efficient. These thresholds can be used to ensure that the retina is never electrically stimulated with currents that lead to off-array activation, a crucial step towards future high-resolution epiretinal implants.

### Algorithm

The algorithm takes as input the electrical activity recorded on the MEA resulting from repeated trials of single-electrode stimulation and determines the lowest stimulus current amplitude at which off-array evoked activity is observed. The voltage waveform recorded on a MEA from epiretinal stimulation (*y)* can be expressed as the sum of four components: *y* = *x* + *i* + *s* + *n*, where *x* is the component resulting from neural activity caused by electrical stimulation, *i* is the electrical voltage artifact resulting from injecting current, *s* is the component resulting from spontaneous (non-evoked) activity of RGCs, and *n* is the noise (such as other biological signals or thermal noise in the recording circuitry). Though *x* is the component that contains information relevant for determining bundle threshold, the other components of y complicate the problem: *i* can overshadow the recorded signal significantly and has properties that are idiosyncratic to the stimulation and recording hardware [5,30], *s* can be confused with activity evoked by stimulation, and *n* can significantly corrupt recorded axonal waveforms. The algorithm overcomes these hurdles by first distinguishing the electrodes recording electrically-evoked spikes from those recording non-evoked spikes or noise by using several characteristic features of extracellularly recorded spikes, followed by identification of bidirectional axonal spike propagation to determine the axon bundle threshold. Three key ideas behind the algorithm are summarized below (and see Figure 2).

**Figure 2.**
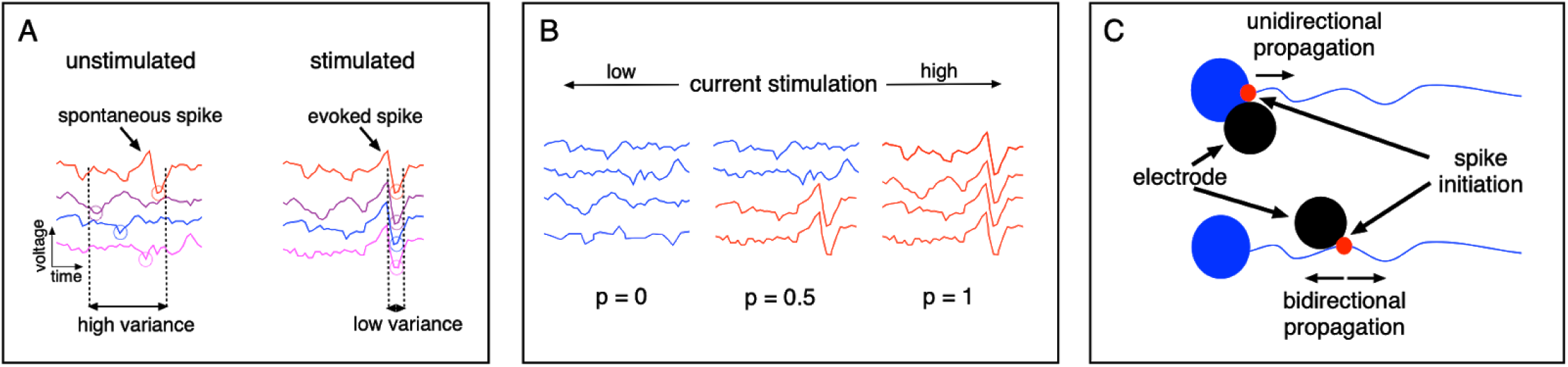
Schematic of ideas exploited in the algorithm. A. Illustration of low-variance in electrically evoked spikes compared to spontaneous spikes. Different traces in a column correspond to recorded traces on an electrode after repeated application of the same electrical stimulus. B. Monotonic increase in response probability as a function of increasing current stimulation [6]. C. Bidirectional axonal spike versus unidirectional somatic spike.

### Idea 1: Precise spike timing

Electrodes recording evoked axonal activity can be distinguished from those recording noise and non-evoked spontaneous activity by leveraging the low variability in the timing of electrically-evoked spikes across repeated trials (Figure 2A). Spike times are estimated by measuring the difference between the time of the stimulation and the time of the minimum response recorded on an electrode. RGC spikes evoked by electrical stimulation occur at a consistent latency, with ∼ 0.1 ms variability between repeats [5–7]. Thus, even if the amplitude of evoked axonal spikes is low, the low timing variability can aid in their detection.

### Idea 2: Monotonic RGC response with stimulation current

The procedure described in idea 1 can still lead to mis-identification of some electrodes recording spontaneous activity as recording electrically-evoked activity. To filter out these electrodes, a second observation is exploited: with increasing stimulus amplitude (in the range 0.1-4.1 μA), spikes are evoked in RGCs with increasing probability and temporal regularity [6]. Thus, if an electrode reliably records an evoked spike at a given amplitude, it will most likely also register that spike upon application of higher current amplitudes, and the spike time variance will not increase (Figure 2B). This allows for more accurate identification of the subset of electrodes recording evoked neuronal activity.

### Idea 3: Bidirectional axonal spike propagation

Finally, the algorithm takes advantage of the fact that axon bundles in any given region of the peripheral retina run approximately in straight lines to the optic nerve, and that signals evoked in stimulated axon bundles travel bidirectionally to opposite edges of the array (Figure 2C). Thus, the axon bundle threshold can be determined by identifying the smallest stimulus current amplitude at which the subset of electrodes recording evoked neuronal activity contains electrodes from at least two borders of the MEA.

A mathematical formulation of the algorithm is provided below. The recorded data consists of a spatio-temporal voltage waveform recorded from the retina using the MEA and is collected after repeated application of single-electrode stimulation, for a range of current amplitudes. Let *e*_*s*_ = stimulating electrode, *e*_*r*_ = recording electrode, *a* = stimulating amplitude, *t* = time (or sample number) after stimulation, *r* = repeat number, 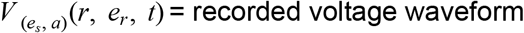 recorded voltage waveform. The algorithm, which can be applied independently on all stimulating electrodes, is described in pseudocode with the steps described more fully below:

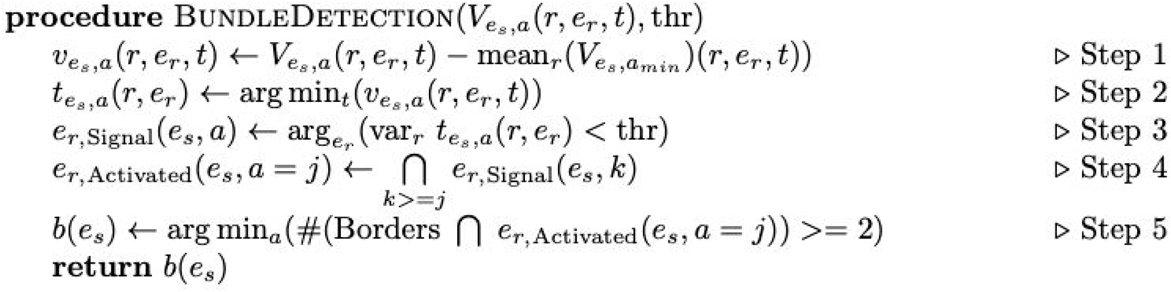

**Algorithm Pseudocode**. The algorithm takes electrical activity recorded on the MEA resulting from repeated trials of single-electrode stimulation as input 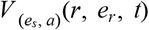 to determine the lowest stimulus current amplitude at which any off-array activity is observed *b*(*e*_*s*_). Statistical threshold based on p-value (thr) is the only hyperparameter in the algorithm.

#### 1. Subtract Electrical Artifact

The recorded signal includes an electrical artifact resulting from the charge supplied during stimulation. To reduce the effect of this artifact on axon bundle threshold detection, the mean recorded data at the lowest stimulation amplitude is subtracted from the recorded data at higher stimulation amplitudes (note that scaling the estimated artifact according to stimulation amplitude did not influence the results). This is repeated for every pair of stimulating and recording electrodes. The voltage trace after artifact removal is referred to as 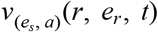.

#### 2. Extract Spike Times

Next, the spike times 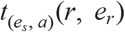 are estimated by measuring the difference between the time of the stimulation and the time of the minimum response recorded on an electrode (the minimum is used because spikes have negative peaks when recorded extracellularly [4,5,31]). Spike times are extracted on each recording electrode after stimulation at one electrode, for every current amplitude and trial. Only the voltage recorded between 0.3−2 ms after the stimulus is considered for spike time extraction, in order to a) avoid large initial recorded artifact and peak variation in artifact across amplitudes recorded at non-stimulated electrodes [32], and b) account for evoked spike latencies and axonal spike propagation [4,6].

#### 3. Extract Signal Electrodes

When no electrically-evoked activity is recorded, the neural response is random in time with respect to the applied stimulus. Thus, the variance of 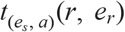 is modeled as a χ^2^_(*n*−1)_ distribution under the assumption that 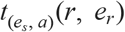 is uniformly distributed across all possible time samples. Here, *n* is the number of repeats and *n* = 25 in the present data. The recording electrodes carrying electrically-evoked activity are then extracted by a hypothesis test with a p-value of 0.05. This statistical threshold based on p-value of the hypothesis testing is the only hyperparameter in the algorithm.

#### 4. Prune Signal Electrodes

Some of the electrodes not recording evoked spikes may also be inadvertently included in the above step. Thus, a pruned set of electrodes at each stimulation amplitude *a* (for each *e*_*s*_) is calculated by finding the common signal-carrying electrodes between this amplitude and all higher amplitudes, to enforce that the electrodes identified as carrying a bundle signal exhibit monotonicity of response with current amplitude (Figure 2B). For each stimulating electrode, pruning is an iterative process that starts at the highest stimulation amplitude and, after calculating the signal-carrying electrodes, works its way down to lower amplitudes.

#### 5. Determine Axon Bundle Threshold

Finally, for each stimulating electrode the current amplitude at which the set of pruned electrodes contains electrodes situated at more than one border of the rectangular MEA is identified as the axon bundle threshold.

## Results

The algorithm was applied to detect bundle activation in *ex vivo* electrical stimulation and recording data from peripheral macaque retina. The results indicate that the algorithm is able to accurately and efficiently detect axon bundle activation while being robust to the selection of the sole hyperparameter (statistical threshold).

### Experimental Setup

Electrophysiology data were collected from retinas of terminally anesthetized rhesus macaque monkeys (male and female, ages 11–20 years), which were euthanized in the course of experiments in other laboratories. Segments of peripheral retina were isolated and mounted on an array of extracellular microelectrodes as described in previous studies [4–6,32,33]. A custom multi-electrode system was used for stimulation and recording spikes in RGCs [5,13,31]. The MEA consisted of 512 electrodes with 60 μm spacing between electrodes, within rows and between rows. For recording, raw voltage signals were amplified, filtered (43-5000 Hz), multiplexed with custom circuitry, and sampled at 20 kHz per channel. For stimulation, charge-balanced triphasic current pulses with 50 μs phase widths and relative phase amplitudes of 2:-3:1, were delivered through one electrode at a time [6]. Custom circuitry included in the stimulation and recording system minimized electrical artifacts, permitting detection of low-latency (<1 ms) RGC responses [6,31]. All reported current amplitudes and polarities refer to the second phase of the pulse, with positive values indicating cathodic currents. In single-electrode scans, each electrode was stimulated repeatedly 25 times with this pulse at each of 40 current levels, progressively increasing by 10% in amplitude over the range 0.1-4.1 μA.

Figure 3 illustrates the algorithm applied to the recorded retinal data.

**Figure 3.**
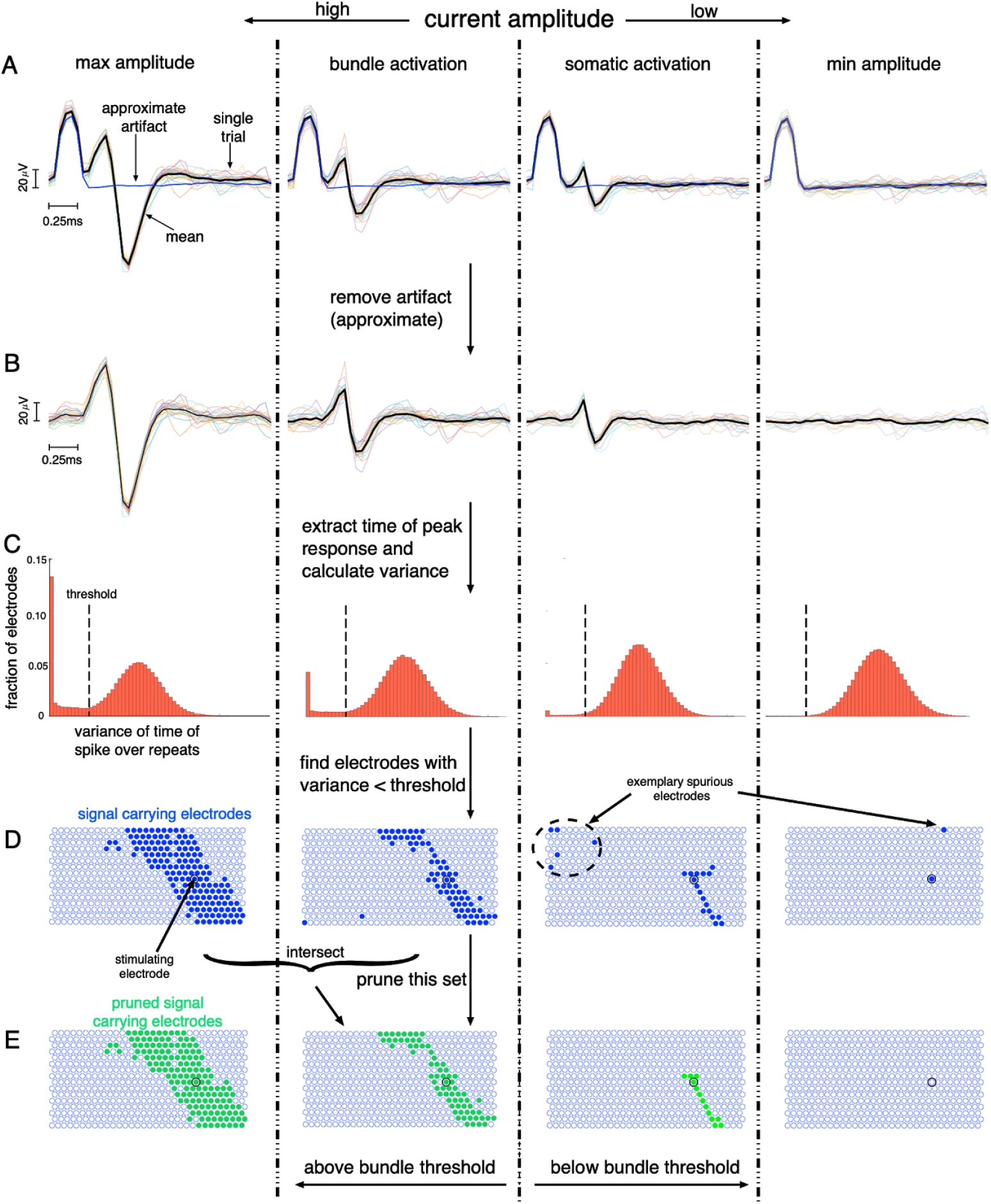
Application of the algorithm to MEA recording and stimulation from the retina. *Left to Right*: Stimulation with decreasing current amplitude showing cases for several amplitudes: highest, amplitude eliciting axon bundle activation, amplitude eliciting somatic spike, and lowest. Top to Bottom: Example cases observed during execution of the algorithm. A. Raw data traces for a particular stimulating and recording electrode. Black trace shows the mean recorded voltage and colored traces show recorded voltage for individual repeats. Blue trace shows the estimated artifact. B. Data traces for the same stimulating and recording electrode after estimated artifact removal. C. Histogram over all recording electrodes of variance over repeats in the time of the spike, for the stimulating electrode chosen above. Dashed vertical line shows the variance threshold below which the recording electrode potentially contained electrically-evoked activity. D. Extracted signal-carrying electrodes shown as mapped onto the MEA. The stimulated electrode is shown as a black ring. The right two panels show clear spurious signal electrodes - electrodes far from the stimulating electrode are not expected to carry electrically-evoked activity. E. Pruned set of signal electrodes and the detection of bundle threshold for the chosen stimulating electrode. The right two panels show the removal of distant spurious recording electrodes.

### Validation against Manual Analysis

To demonstrate the effectiveness of the algorithm, automatically estimated axon bundle thresholds were compared to values estimated using manual analysis by experienced researchers observing movie clips of evoked electrical activity. A total of ∼1,500 stimulating electrodes from four different retinal preparations were analyzed. In the movie clips, the electrical artifact was reduced by subtracting the mean activity recorded at the lowest stimulation amplitude from the traces recorded with all higher stimulation amplitudes. The result was then averaged over multiple trials of stimulation at each amplitude. Recorded voltage amplitudes were max-thresholded at 36 μV to help facilitate tracking of low amplitude axonal activity in the midst of high-amplitude somatic activity. This max-thresholded spatio-temporal activity, 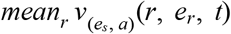, was viewed for each stimulation electrode and amplitude. Observers estimated the lowest stimulus amplitude required to evoke bidirectional electrical activity propagating all the way to the edges of the MEA. For some of the stimulating electrodes on or near the border of the MEA, only a few recording electrodes were able to capture evoked activity. This fact, combined with large residual stimulation artifacts in nearby recording electrodes, made manual determination of bidirectional activity difficult in these cases. Stimulating electrodes in these cases were not assigned a manual bundle threshold and were not used in algorithm validation.

The thresholds identified by the algorithm were similar to those identified by a trained human observer, with a correlation coefficient of 0.95 (Figure 4B). For ∼88% of the electrodes analyzed, the threshold identified by the algorithm was within ±10% of the manually identified threshold (corresponding to ±1 amplitude step in the stimulation experiment). For 65% of electrodes, the match was exact (Figure 4A). To compute the probability of obtained thresholds as compared to chance, manual thresholds for the electrodes of each retina were randomly permuted and then pooled. In these permutations, only 24 ± 1% of the algorithm thresholds were within 10% of the manual thresholds, a substantially lower fraction than observed in the data. To quantify the reliability of the manually labeled data, the intrinsic variability between two human observers who were asked to perform the same task was examined. For stimulating electrodes at which both humans assigned a threshold, ∼96% of the electrodes had a threshold within ±10% of each other, with a correlation of 0.98, similar to the values obtained by comparing the manual and algorithm thresholds.

**Figure 4.**
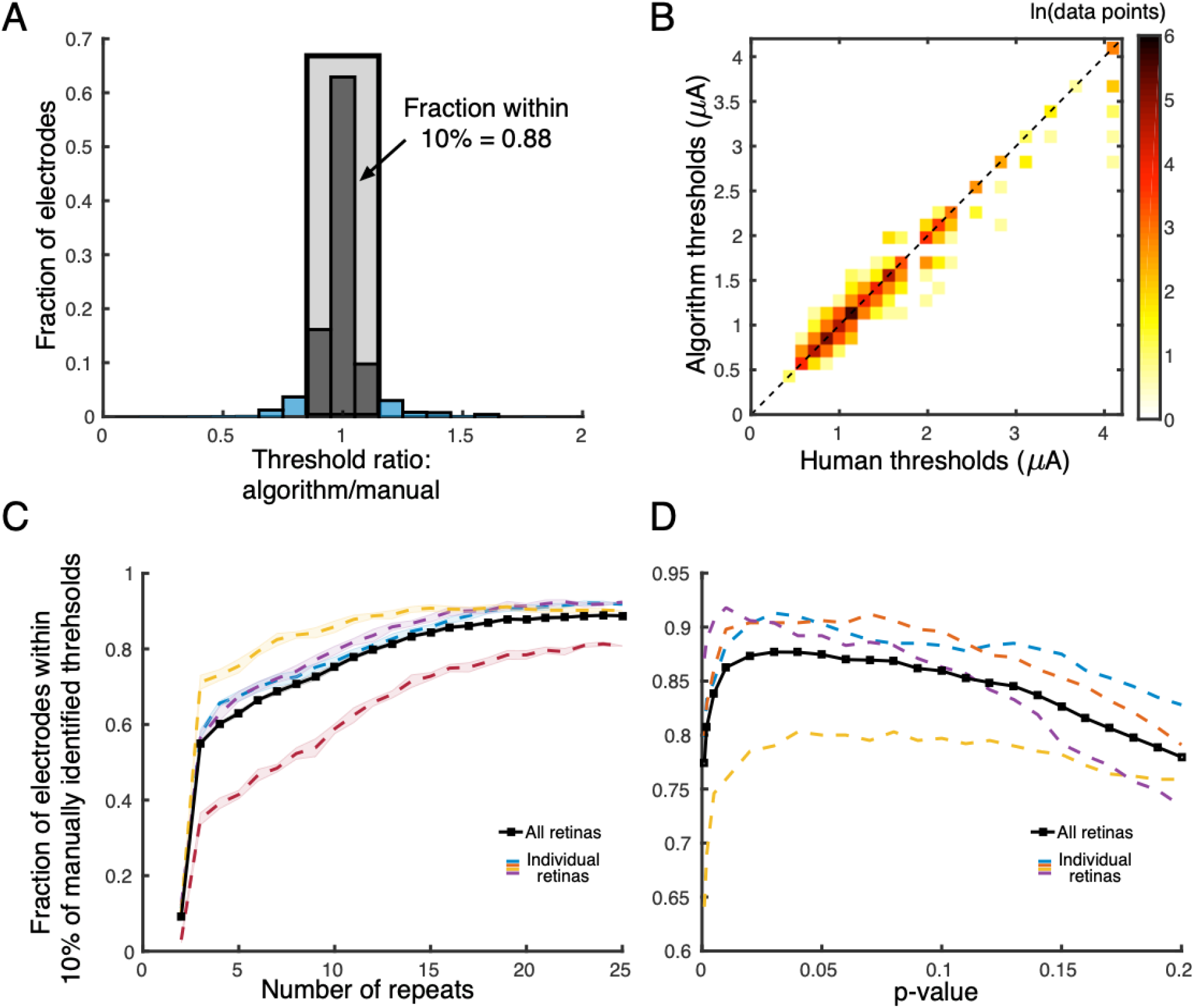
Validation against manual analysis. A. Histogram over stimulating electrodes of ratio between algorithm and manual axon bundle thresholds. ∼88% of the electrodes analyzed exhibited an algorithmic axon bundle threshold within ±10% of the manually identified threshold, for four different retina preparations (∼1500 stimulating electrodes). B. Scatter plot of algorithm thresholds and manually analyzed thresholds. Color represents the density of points with a particular set of thresholds (note that there are 40 discrete current stimulations in the data). The clustering of the data around the diagonal of equality suggests that the algorithm is not biased (correlation coefficient = 0.95). C. Dependence of the accuracy of the algorithm (compared to manually identified thresholds) on the number of repeats. Different colors correspond to four different retinal preparations. Data were randomly subsampled from the maximum available repeats (25), and the shaded region encompasses one standard deviation. The algorithm performance exhibits diminishing returns with increasing repeats, with apparently saturated performance at 25 repeats. D. Algorithm performance as a function of the statistical threshold (p-value). Performance remains consistent around the chosen p-value of 0.05.

### Further observations

#### The algorithm is efficient and performs well with limited electrophysiological data

Because collecting and analyzing a large amount of electrical stimulation data is difficult, it is advantageous to limit the number of stimulation repeats required for the algorithm to perform well. To determine the data requirements, comparison of automatically determined thresholds to manually determined thresholds was performed after running the algorithm on a random subset of the original 25 stimulation trials. Manually identified thresholds were obtained using all trials. The performance of the algorithm followed a saturating curve with accuracy increasing sharply from 1-3 repeats and leveling off with 20-25 repeats for all retinas (Figure 4C). On average, the algorithm identified the bundle threshold on ∼84% electrodes within ±10% of the manually identified threshold with only 15 repeats and ∼87% with only 20 repeats. However, for some retinas, having access to more stimulation repeats was advantageous (Figure 4C, red curve).

#### The algorithm performance is robust to the sole hyperparameter

To test whether the observed results also generalize to new retinas, variation in the performance of the algorithm was studied across different retinal preparations and values of the statistical threshold (p-value) used for hypothesis testing. The p-value is the only design hyperparameter in the algorithm (used for finding the subset of electrodes recording evoked activity; see Algorithm) and could lead to a suboptimal algorithm performance if not chosen appropriately. The average performance of the algorithm was within ±0.5% for a range of thresholds corresponding to the p-value range of 0.02-0.08 (Figure 4D). This robust behavior is likely due to the strong monotonic response requirement immediately after the hypothesis testing step (see Algorithm). Therefore, the threshold corresponding to the p-value of 0.05 was used for all retinas in the reported results, irrespective of its optimality for individual retinas.

### Conclusions

This work presents an automated approach to detect activation of off-array cells via their axons, known as axon bundle activation, using stimulation and recording data from a MEA. The algorithm consists of simple computational motifs, and could be implemented easily on hardware for continuous monitoring of axon bundle thresholds in an *in vivo* implant. Implementation within power and thermal constraints is an important future research topic. The algorithm can be used in a closed-loop fashion by a future epiretinal prostheses to rapidly determine the axon bundle thresholds during calibration. Though the focus of this work was on epiretinal prosthesis, the need to detect activation of distant cells via their axons will exist in most high-resolution prosthetic systems, and the algorithm has the potential to generalize to such systems.

## Acknowledgements

Stanford Graduate Fellowship (PT), Wu Tsai Neurosciences Institute Big Ideas (EJC, SMi), NIH NEI R01 EY021271 (EJC), Polish National Science Centre Grant DEC-2013/10/M/NZ4/00268 (PH).

